# Tracking gene expression of single mitochondria in live neurons using nanotweezers

**DOI:** 10.64898/2025.12.10.693488

**Authors:** Annie Sahota, Binoy Paulose Nadappuram, Siân C. Allerton, Flavie Lesept, Jack H. Howden, Suzanne Claxton, Yaxian Liu, Francesco A. Aprile, Josef T. Kittler, Michael J. Devine, Aleksandar P. Ivanov, Joshua B. Edel

## Abstract

Neurons are highly polarised cells that depend on mitochondria for energy and signalling homeostasis. Importantly, energy and signalling requirements vary considerably across individual neurons both spatially and temporally. Therefore, to fully understand neuronal mitochondria, methods are needed to analyse mitochondria in live cells over time. The nanotweezer, a minimally invasive single-cell sampling technique, enables precise extraction a individual mitochondria from defined subcellular locations. Here, we combine single-mitochondrial extraction from live neurons with mitochondrial gene expression tracking and mtDNA profiling. By tracking mitochondrial gene expression in the same neurons over time, we reveal a downregulation of mitochondrial genes MT-ND1 and MT-ATP6 following exposure to α-synuclein aggregates, independent of the proximity of the aggregates to the sampled mitochondria. Our approach provides precise, dynamic measurements of mitochondrial composition and gene expression in vivo at single-organelle resolution, enabling mechanistic studies of neuronal mitochondrial heterogeneity and its perturbation in models of neurodegeneration.

## Main

Neurons rely heavily on mitochondria because of their disproportionately high ATP demand compared to other cell types.^1^ Mitochondrial dysfunction is a core feature of ageing and neurodegeneration, including Parkinson’s disease (PD),^2^ Alzheimer’s disease (AD)^3^ and amyotrophic lateral sclerosis (ALS).^4^ Decreased mitochondrial DNA (mtDNA) copy number and elevated mtDNA heteroplasmy have been identified in neurodegenerative diseases.^5^ Dysregulation of mitochondrial gene expression and mitochondrial dynamics has also been observed, affecting synaptic transmission, membrane potential maintenance, calcium buffering, and apoptosis.^6^

Mitochondrial subpopulations are evident within neuronal subcompartments.^7^ For example, synaptic mitochondria are distinct in their morphology,^8,9^ proteomic profiles,^10^ enzyme activity,^11^ calcium buffering^12^ and vulnerability compared to non-synaptic mitochondria.^13^ This, combined with the knowledge that specific compartments of a neuron may be selectively degenerated in neurodegenerative disease before the loss of the neuron,^14,15^ highlights the need to study the heterogeneity of compartmentalised mitochondria within single neurons. Single-cell analysis technologies have enabled the characterisation of mitochondrial heteroplasmy in neurons and hold great potential for a better understanding of heteroplasmy at the single-cell level.^16,17^ For example, mitochondrial transplantation has been performed using FluidFM, in which multiple mitochondria and portions of the mitochondrial network were transplanted from HeLa cells into U2OS cells and primary human endothelial keratinocytes.^18^ More recently, an automated nanobiopsy platform has been developed for mitochondrial transplantation and sensing.^19^

Understanding the spatial differences in mitochondrial distribution in neurons and the localised effects of pathological markers, such as protein aggregation, on neuronal mitochondria could be further advanced by analysing the composition of individual mitochondria in live neurons. This could enable precise mitochondrial mechanisms to be unravelled and novel treatment targets to be identified at single organelle resolution. Steps towards this have been shown in a recent high-resolution imaging study, in which mitochondrial translation products were observed in the axons, dendrites, and synaptic regions of hippocampal neurons.^20^ Additionally, single-cell sampling of individual mitochondria from neurons using micropipettes and nanopipettes has been combined with sequencing to study mitochondrial heteroplasmy.^21–23^ However, no method currently enables the isolation and analysis of individual mitochondria for tracking mitochondrial gene expression in live cells. Further advances in single-cell sampling approaches and analysis methods to examine the composition of individual mitochondria in neurons have the potential to improve our understanding of basic mitochondrial biology in neurons substantially and to facilitate the discovery of novel cellular mechanisms and the development of improved therapeutic interventions against neurodegenerative diseases.^24^

To bridge this gap, we build on our nanotweezer technology, which enables minimally invasive biomolecular extraction from living cells. We previously demonstrated single-mitochondrion extraction from neuronal axons and repeated cytoplasmic sampling without compromising viability. ^25–27^ More recently, we have shown that multiple cytoplasmic extractions can be performed on the same live neuron, including from neurites, without affecting cell viability, a feat particularly notable given the delicate nature of neurons *in vitro*.^27^ Here, we develop an approach that combines multiple individual mitochondrial extractions from the same neuron with single-mitochondrial mtDNA and mtRNA analysis, enabling profiling of the composition of single organelles in live neurons. Furthermore, we show that individual mitochondria can be isolated from specific neuronal locations and at different time points within the same cell following uptake of α-synuclein (α-syn) oligomers, thereby enabling tracking of mitochondrially encoded gene expression in live neurons in a model of neurodegeneration.

## Results

### Extraction of single mitochondria from live neurons

To isolate individual mitochondria from live neurons, mitochondria in neurites were targeted for nanobiopsy extraction using the nanotweezer (Fig. 1a). Mitochondria were fluorescently labelled *in vitro* by transfecting neurons with mtDsRed (Fig. 1b), which enabled specific targeting of a mitochondrion’s matrix. Using bright-field imaging (Fig. 1b, inset), the nanotweezer was manually inserted into a neuron adjacent to a labelled mitochondrion. The extraction of a single mitochondrion was then monitored by fluorescence following nanotweezer insertion into the cell (Fig. 1c(i), Fig. 1d(i), Fig. 1e(i)). An AC voltage was applied adjacent to a mitochondrion, which enabled the mitochondrion to become trapped at the nanotweezer tip by dielectrophoresis (DEP) (Fig. 1c(ii), Fig. 1d (ii)), illustrated by a shift in mitochondrial fluorescence closer to the nanotweezer tip (Fig. 1e(ii)). The nanotweezer was then removed from the cell (Fig. 1), extracting the trapped mitochondrion, as illustrated by a loss of fluorescence derived from the isolated mitochondrion (Fig. 1d(iii), Fig. 1e(iii)). To further confirm successful mitochondrial extraction from the cell, fluorescence was observed on the nanotweezer tip outside the cell, which persisted when moving the nanotweezer, confirming that the mitochondrion was trapped on the nanotweezer tip.

**Fig. 1.**
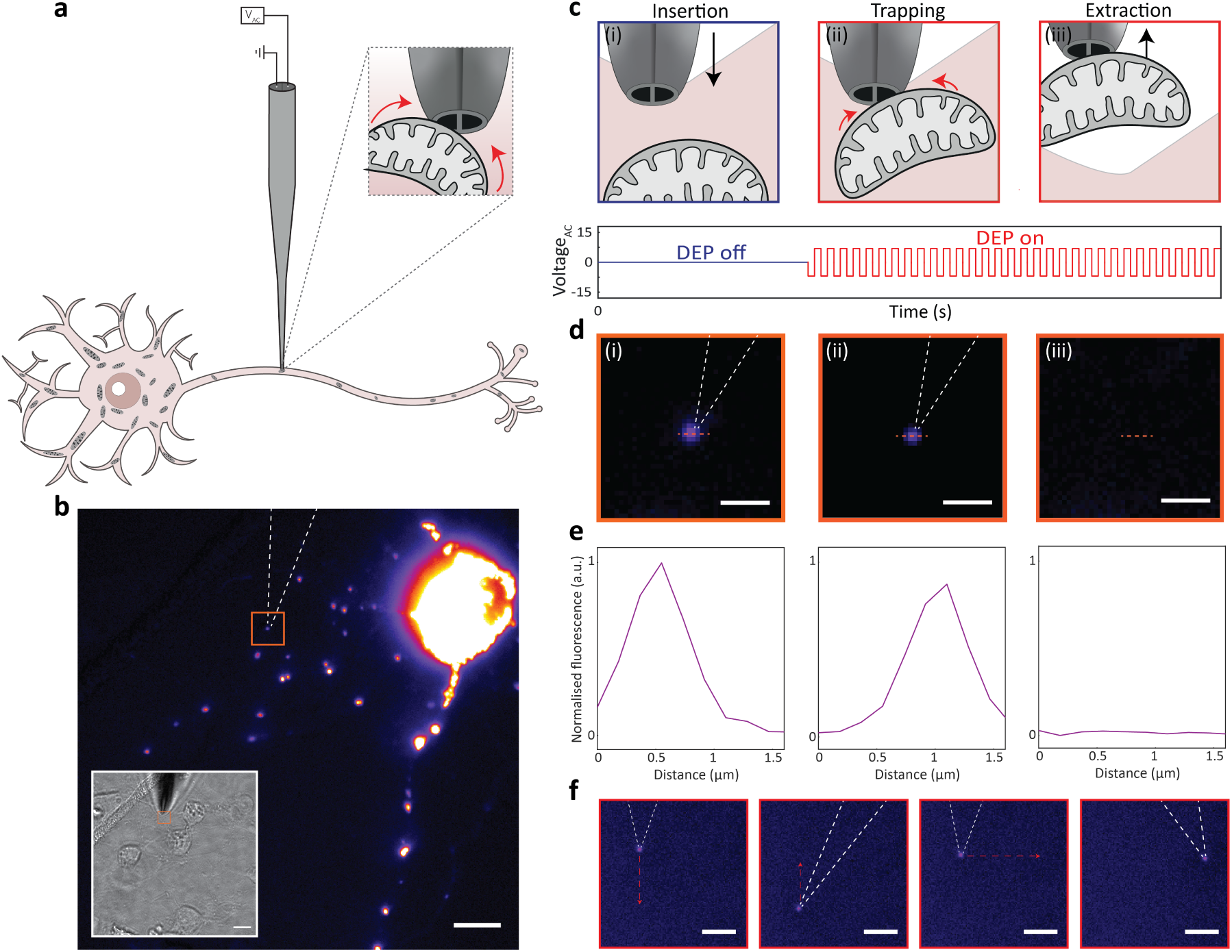
Single mitochondrion isolation. a) Schematic illustrating the trapping of a single mitochondrion from a neuron using the nanotweezer. b) Fluorescent image of an mtDsRed-labelled neuron and corresponding bright field image (inset) of the nanotweezer inserted into the cell adjacent to a single mitochondrion within the cell. The orange box represents the region containing the targeted mitochondrion. The dashed white lines on the fluorescent image represent the position of the nanotweezer. Scale bars = 10 μm. c) Schematic showing insertion of the nanotweezer into the cell adjacent to a mitochondrion (i), DEP trapping of the mitochondrion via application of an AC voltage (ii) and extraction of the mitochondrion via removal of the nanotweezer from the cell. d) Corresponding fluorescent images of the targeted mitochondrion after insertion of the nanotweezer (i), DEP trapping at the nanotweezer tip (ii) and removal of the nanotweezer from the cell (iii), showing complete loss of the targeted mitochondrion’s fluorescence. Scale bars = 2 μm. e) Corresponding fluorescence traces of the targeted mitochondrion measured over the distance shown by the dashed line in (d), showing levels of mitochondrial fluorescence before application of DEP (i) trapping of the mitochondrion at the tip upon application of DEP (ii) and loss of mitochondrial fluorescence following removal of the trapped mitochondrion. f) Fluorescent images of the nanotweezer tip outside the cell after removal of the trapped mitochondrion, where the nanotweezer was moved in x and y directions, and the fluorescent mitochondrion moved simultaneously, confirming the presence of a trapped mitochondrion at the tip. Scale bars = 5 μm.

To assess the viability of the mitochondrion trapped using the nanotweezer and to confirm that intact mitochondria were isolated from cells, a single mitochondrial nanobiopsy was performed with mitochondria labelled with TMRM, a fluorescent probe readily sequestered by mitochondria with intact mitochondrial membrane potential (MMP) (Fig. S1). The fluorescence of the mitochondrion was observed before and during nanotweezer DEP application, as well as after mitochondrial extraction from the cell, where the nanotweezer was maintained in the cell’s culture medium (Fig. S1a). Mitochondrial extraction was confirmed as before, where an increase in fluorescence was observed at the nanotweezer tip upon application of DEP, showing that the targeted mitochondrion moved towards the nanotweezer tip (Fig. S1b). The viability of isolated mitochondria was also validated, with no loss of mitochondrial fluorescence observed during the trapping procedure, and fluorescence remaining on the nanotweezer tip after extraction from the cell (Fig. S1c). To investigate possible effects on the mitochondria that remained in the cell following nanobiopsy, the fluorescence of neighbouring mitochondria and non-neighbouring mitochondria in the same cell was measured during the nanobiopsy procedure. This was compared to the fluorescence of mitochondria in neighbouring cells that were not subject to nanobiopsy. As illustrated by Fig. S1d, the fluorescence intensity of the mitochondria remained constant, showing that there were minimal effects on mitochondrial membrane potential and that the viability of mitochondria in the same cell was not affected. The long-term viability of extracted mitochondria was investigated by measuring the fluorescence intensity of the trapped mitochondria on the nanotweezer tip outside the cell, demonstrating that mitochondria isolated using the nanotweezer retained MMP for at least 90 minutes outside the cell.

### Single mitochondrion analysis

Having demonstrated that intact mitochondria can be extracted from neurons using the nanotweezer without impacting mitochondrial viability, protocols to analyse the content of a single mitochondrion were developed. First, mtDNA was successfully detected using qPCR following single mitochondrial lysis and purification (Fig. 2a). mtDNA was detected exclusively in nanobiopsy samples, as evidenced by the absence of amplification in qPCR negative and purification controls. The negative control was a no-template control that comprised all the qPCR reagents, along with nuclease-free water, instead of the DNA. The purification control comprised the lysis and purification steps applied to nuclease-free water. To investigate potential non-specific adherence of mitochondria or mtDNA to the nanotweezer, the nanotweezer was inserted into the cell’s media and into the cell itself without the application of DEP. These additional controls showed minimal amplification compared to the amplification of extracted mitochondria. Following the detection of mtDNA from each mitochondrion, mtDNA copy numbers from individual mitochondria were obtained (Fig. 2b, Fig. S2). Amplification was observed in 9 out of 14 mitochondria extracted using the nanotweezer, with copy numbers between 0 and 13 per mitochondrion. These copy numbers are similar to the previously reported estimates of mtDNA copies per mitochondrion.^28,29^ It is important to highlight that absolute qPCR quantification is based on the construction of a standard curve where, despite the standard curve constructed in this work having a high R^2^ value (0.99) and the primers having good PCR efficiency (96%), slight variations from pipetting, amplification efficiencies per well, and sample mixing may affect the quantification, particularly when assessing copy numbers in the range described here. Single mitochondrial mtDNA quantification used in this manner may therefore be suited for relative comparisons between samples processed in parallel rather than for examining absolute numbers within a single sample type. For example, the method described here could be used to compare mtDNA copy numbers within subcellular compartments of the same neuron, as has been recently described.^30^ Alternatively, the method could be used to compare mitochondria in healthy and PD neurons.

**Fig. 2.**
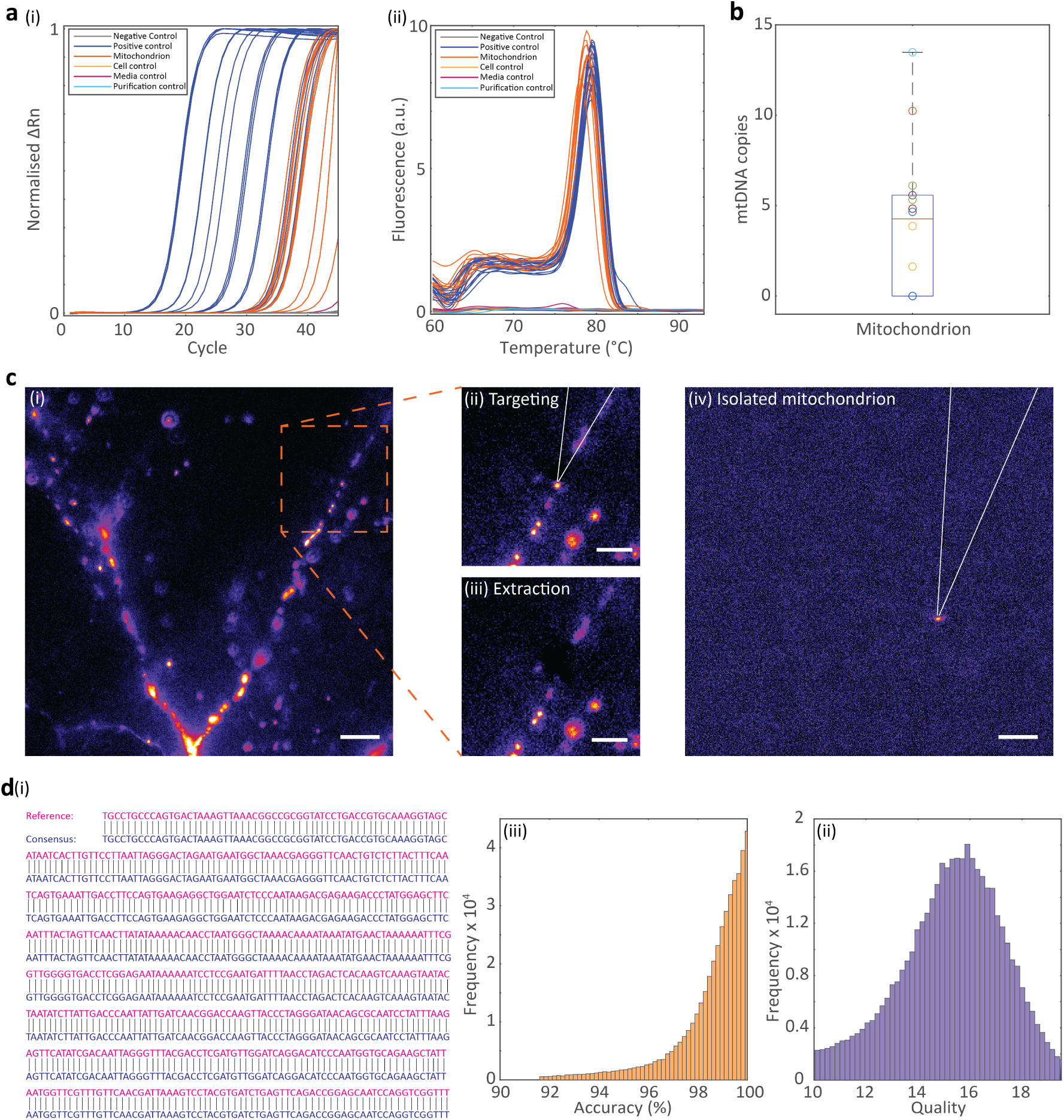
Single mitochondrion mitochondrial DNA analysis. a) qPCR amplification plot (i) and melt curve (ii) of mtDNA isolated from individual mitochondria extracted from neurons compared to controls. qPCR was performed using primers for MT-CO1. b) mtDNA copy numbers calculated for each mitochondrion (n=14 mitochondria). c) Fluorescent images of mtDsRed-transfected neurons and targeted mitochondrion for extraction and sequencing. The dashed box in (i) represents the targeted region in the cell. The nanotweezer was targeted to an individual mitochondrion in this region (ii), followed by extracting the mitochondrion (iii). Isolation of the mitochondrion was confirmed by a loss of fluorescence in the cell following extraction and the detection of fluorescence at the nanotweezer tip outside the cell (iv). The white lines represent the nanotweezer. (i) and (iv) scale bars = 10 μm. (ii) and (iii) scale bars = 5 μm. Summary statistics for boxplots: centre = median; bounds of box = IQR 25th and 75th percentile; whiskers = minimum and maximum within 1.5 IQR. d) Overall alignment (i), accuracy distribution (ii) and quality distribution (iii) of the reads from nanopore sequencing of a 558 bp region of the mtDNA from the extracted mitochondrion.

Mitochondria extracted using the nanotweezer were then analysed by DNA nanopore sequencing. Single-mitochondrial sequencing without amplification was unsuccessful, likely due to the small number of mtDNA copies per mitochondrion and the potential loss of material during library preparation. To address this, mitochondrial DNA was sequenced from PCR-amplified samples. A single mitochondrion was isolated from a neuron using the nanotweezer (Fig. 2c), lysed, purified, and then a 558 bp region within the MT-RNR2 gene was amplified using a high-fidelity polymerase. PCR-amplified material was then successfully sequenced by nanopore sequencing, as demonstrated by 100 % alignment of the consensus sequence with the reference sequence (Fig. 2d(i)), a median quality score of 15.0 (Fig. 2d(ii)), and a median accuracy score of 98.4% (Fig. 2d(iii)).

A protocol for analysing RNA from a single mitochondrion was also developed. As the amount of mtDNA in each mitochondrion can differ, exclusion of mtDNA is required for gene expression analysis. Due to mitochondrially encoded RNAs not containing introns, this could not be performed using primers that spanned exon-exon junctions in RT-qPCR. To address this, the purification step following mitochondrial lysis was modified by utilising surface-functionalised magnetic beads with target-specific binding regions. The target-specific binding regions comprised oligonucleotides with a repeated sequence of 25 thymine nucleotides for binding to polyadenylated RNAs and oligonucleotides with a sequence complementary to 12S rRNA. This enabled the selective enrichment of mtRNAs during purification and avoided the carryover of mtDNA, which may otherwise lead to false-positive gene expression analysis. The single mitochondrion mtRNA analysis method is summarised in Fig. 3a.

**Fig. 3.**
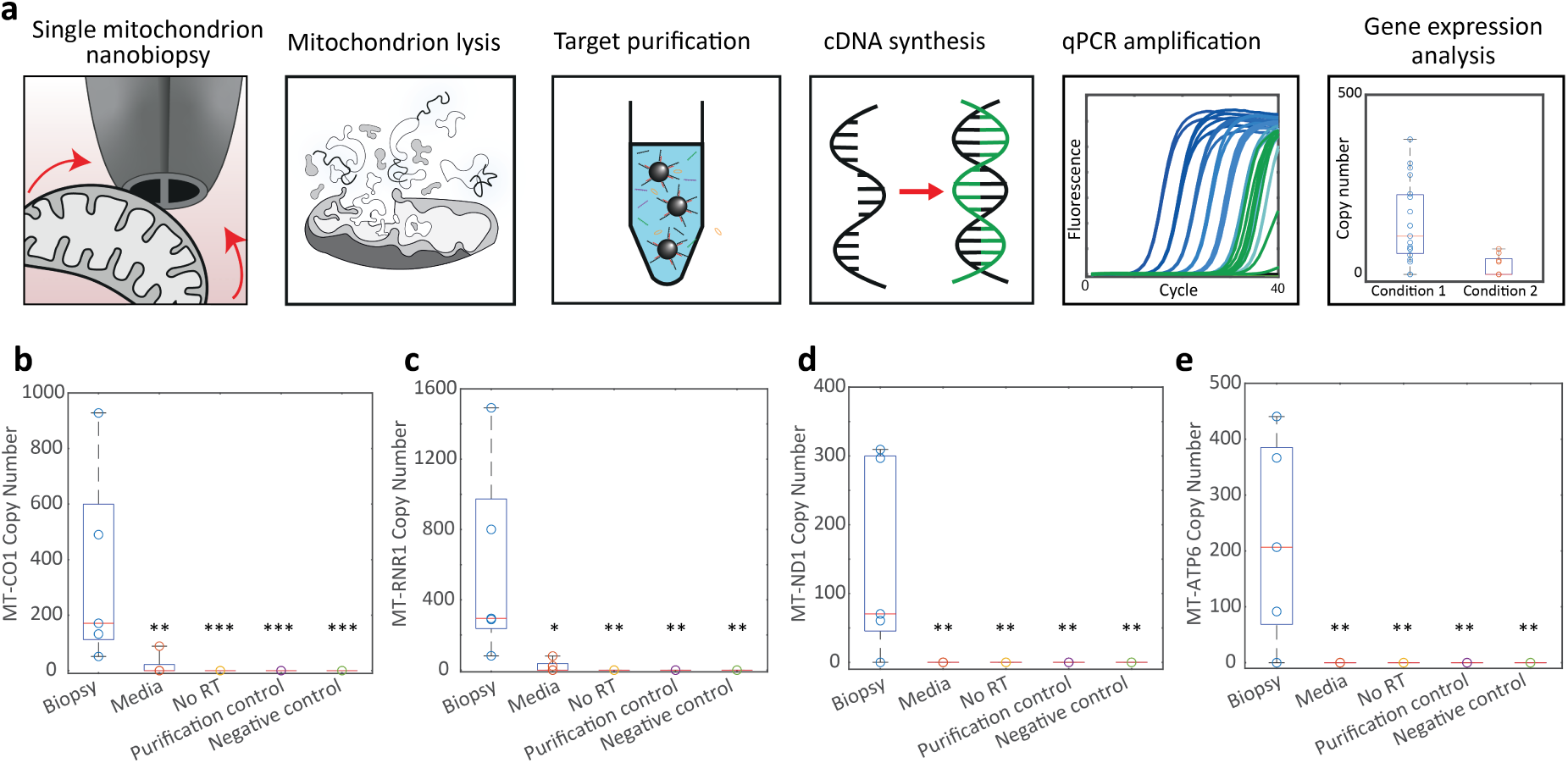
Single mitochondrial RNA analysis. a) Schematic of single mitochondrial mtRNA analysis procedure. b) RT-qPCR copy numbers derived from single mitochondrial nanobiopsies, cell culture media controls, no RT controls, and RT-qPCR negative controls for MT-CO1, c) MT-RNR1, d) MT-ND1 and e) MT-ATP6 expression (n=5). Statistical analysis was performed using the Kruskal-Wallis non-parametric test between biopsies and all controls (*P < 0.05, **P < 0.01, ***P < 0.001, ^n.s^P = not significant). Summary statistics for boxplots: centre = median; bounds of box = IQR 25th and 75th percentile; whiskers = minimum and maximum within 1.5 IQR.

The isolation and detection of mtRNA in the absence of mtDNA were successful, as evidenced by the absence of mtDNA in the no-RT controls. The detection of mtRNA from the nanobiopsy extractions was further supported by the detection of a significantly higher mtRNA concentration in mitochondria isolated by nanobiopsy than in the purification, media and RT-qPCR controls. The purification control served as a further negative control, in which water was analysed using the method shown in Fig. 3a to ensure that the detected mtRNA did not arise from contamination during the analysis. The media control represented mtRNA derived from nanobiopsy from media outside the cell, accounting for any potential cell-free mitochondria or mtRNA. Combined, these results show that mtRNA can be analysed from an individual mitochondrion extracted using the nanotweezer and that the detected mtRNA derived from the extracted mitochondrion.

### Spatially-resolved single mitochondrial extraction shows that proximity of α-synuclein oligomers does not determine mitochondrial gene expression

To demonstrate the capability to probe specific mitochondrial gene expression profiles within a single neuron, the interaction between α-synuclein (α-syn) aggregates and mitochondrial gene expression was investigated. Previous work has suggested a direct interaction between α-syn aggregates and neuronal mitochondria,^31–33^ in which α-syn (including oligomeric species) can pass through cellular membranes and localise within mitochondrial membranes near mitochondrial proteins such as ATP synthase (complex V) and mitochondrial complex I.^34–37^

As small, soluble oligomers are considered the most toxic form of α-syn,^38^ mixtures comprising both oligomers and monomers were prepared. Aggregation and oligomer formation were confirmed by atomic force microscopy (AFM), as evidenced by an increase in equivalent disc radius and a broader size distribution with incubation (Fig. S3). To investigate the effect of α-syn oligomers only, the protein mixtures were labelled with an α-syn aggregate-specific fluorescently labelled aptamer.^39^ The fluorescently labelled α-syn oligomers were incubated with mtDsRed-labelled neurons for 24 h to enable their uptake into cells, and then washed to remove the excess (Fig. 4a). This enabled both the α-syn oligomers and mitochondria to be imaged in the neurons.

**Fig. 4.**
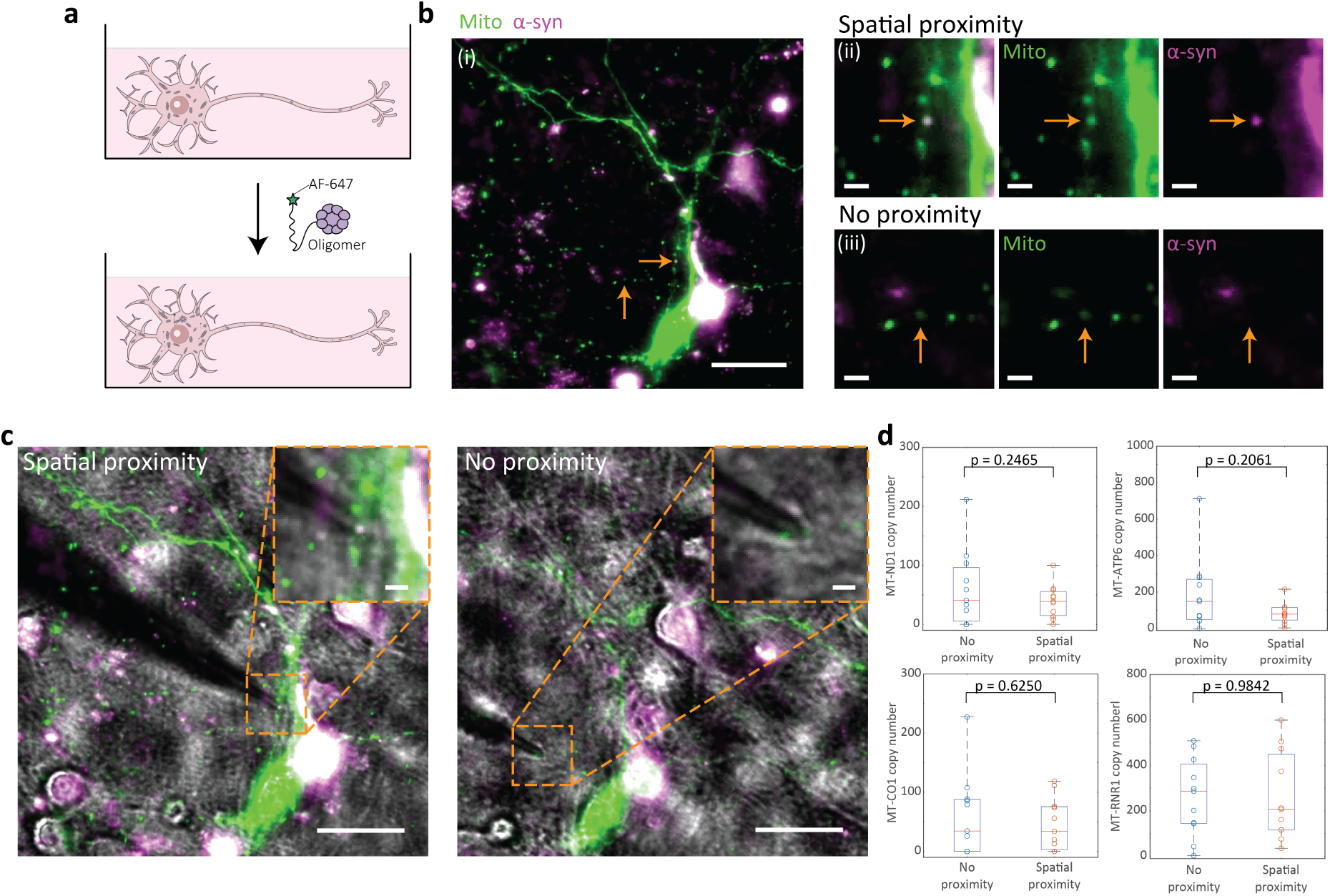
Mitochondrially-encoded gene expression from single mitochondrion biopsies in spatial proximity with *α*-synuclein oligomers. a) Schematic of the application of fluorescently labelled α-synuclein oligomers to neurons. b) Fluorescent images of mtDsRed-labelled DIV 14 neurons following 24 h incubation with fluorescently labelled α-synuclein oligomers (i). Some of the fluorescently labelled α-synuclein oligomers were in spatial proximity to the mitochondria in the cell (ii), and some were not (iii). Mitochondria are labelled in green, and α-synuclein oligomers are labelled in magenta. (i) scale bar = 20 μm, (ii) and (iii) scale bars = 2 μm. c) Fluorescent images of mtDsRed-labelled neurons with fluorescently labelled α-synuclein oligomers overlaid with bright field images of nanobiopsies of single mitochondria in spatial proximity with α-synuclein oligomers and in no proximity with α-synuclein oligomers. Scale bars = 20 μm. Inset scale bars = 2 μm. d) Boxplots of MT-ND1, MT-ATP6, MT-CO1 and MT-RNR1 copy numbers derived from individual biopsies of mitochondria proximal and not proximal to α-synuclein oligomers (n = 11 mitochondria). Statistical analysis was performed using a Wilcoxon test (MT-ATP6, MT-CO1) or a paired t-test (MT-ND1, MT-RNR1). Summary statistics for boxplots: centre = median; bounds of box = IQR 25th and 75th percentile; whiskers = minimum and maximum within 1.5 IQR.

As shown by the overlap between the α-syn channel and the mitochondrial channel in Fig. 4b(i), some of the α-syn oligomers were near individual mitochondria (Fig. 4b(ii)), and some were not (Fig. 4b(iii)) following uptake of the α-syn oligomers in the cells. Spatial proximity was defined by overlap between α-syn and mitochondrial channels, as shown in Fig. 4b(ii). To investigate whether this proximity influenced single mitochondrial gene expression, individual mitochondria overlapping and not overlapping with α-syn oligomers were extracted using the nanotweezer (Fig. 4c). The nanotweezer’s high spatial precision enabled the specific selection of mitochondria. Gene expression from each mitochondrion was analysed using the method outlined in Fig. 3a using standard curves for absolute copy number quantification of each mitochondrion (Fig. S4).

Surprisingly, no significant difference between the expression of MT-ND1, MT-ATP6, MT-CO1 and MT-RNR1 was observed in mitochondria that overlapped with α-syn oligomers and mitochondria that did not. Therefore, these findings do not support a direct influence of α-syn oligomers on mitochondrial gene expression.

### Single-cell mitochondrial gene expression tracking shows suppression of mitochondrially encoded gene expression due to α-syn oligomer exposure

To investigate the effects of α-syn oligomers on mitochondrial gene expression in neurons, the nanotweezer was used to extract multiple individual mitochondria from the same live neuron over 24 h. Single-cell gene expression tracking enables the same live cell to be probed, uncovering changes in gene expression and unmasking the heterogeneity of cell populations that is missed in bulk studies. The nanotweezer was therefore used to examine the potential effects of α-syn oligomers on mitochondrial gene expression at the single-neuron level.

Fig. 5a shows a schematic of the single mitochondrial extraction procedure that was performed in the same cells using the nanotweezer to track mitochondrial expression following the addition of α-syn oligomers. Using gridded cell culture dishes to locate the regions of neurons undergoing nanobiopsy, multiple single mitochondria were extracted from the same cell under normal culture conditions (Fig. 5b, before treatment). Aggregated α-syn mixtures were then incubated with the cells for 24 h, after which the cells were washed to allow α-syn to be taken up. Following this, the same cell was located, and multiple individual mitochondria were extracted from the cell again using the nanotweezer (Fig. 5b, after treatment). This provided populations of individual mitochondria from single cells at two timepoints. The mitochondrial gene expression from each mitochondrion was then analysed using the method shown in Fig. 3a, and overall changes in MT-ND1, MT-ATP6, MT-CO1 and MT-RNR1 expression in the same cells were assessed. This was compared with mitochondria derived from corresponding control cells, in which a buffer containing no protein was applied instead of the α-syn mixture.

**Fig. 5.**
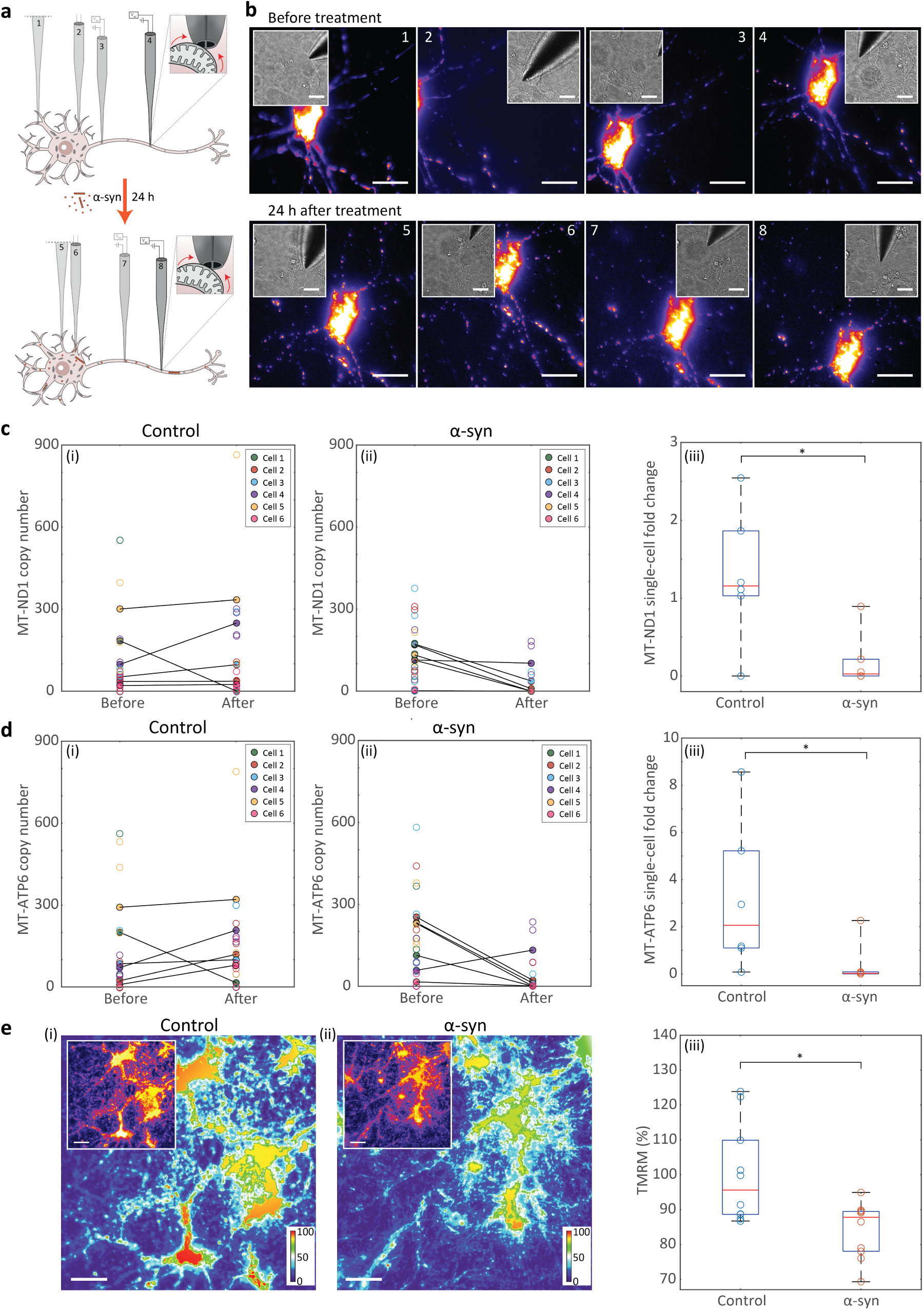
Single-cell tracking of mitochondrially encoded gene expression with incorporation of aggregated *α*-synuclein. a) Schematic of same-cell single mitochondrial biopsies with application of aggregated α-synuclein. 3-4 single mitochondria were extracted from a DIV 14 neuron, 0.5 μM aggregated α-synuclein was incubated with the cells for 24 h, the cells were then washed, and a further 3-4 single mitochondria were extracted from the same cell. b) Representative fluorescent images of 8 nanobiopsies taken from the same cell. Inset: bright field images of the nanotweezer inserted into the cell at the targeted mitochondrion. Scale bars = 20 μm. c) Expression of MT-ND1 tracked in control cells (i), expression of MT-ND1 tracked in α-synuclein-treated neurons (ii), and (iii) the fold change of MT-ND1 measured in the same cells in control and α-synuclein-treated neurons (n=6 cells). d) Expression of MT-ATP6 tracked in control cells (i), expression of MT-ATP6 tracked in α-synuclein-treated neurons (ii), and (iii) the fold change of MT-ATP6 measured in the same single cells in control and α-synuclein-treated neurons (n=6 cells). Statistical analysis was performed using a Mann-Whitney non-parametric test (*P < 0.05, ^n.s^P = not significant). Each filled circle in the tracked plots represents the mean copy number of each cell; the unfilled circles represent the individual copy numbers from each mitochondrion, and the black lines link the mean copy numbers in the same cell before and after treatment. Each unfilled circle in the fold change plots represent the fold change between the mean mitochondrial copy numbers in the same cell before and after treatment. e) Heat map of TMRM-loaded control cells (i), heat map of TMRM-loaded cells following 24 h treatment with aggregated α-synuclein (ii), and normalised TMRM fluorescence of control and aggregated α-synuclein-treated cells measured as a percentage of the mean normalised fluorescence of control cells (n=10). TMRM fluorescence was normalised to the cell area in each image. Colour bars = fluorescent values of the image (arbitrary units). Inset: Corresponding fluorescent images of the TMTM-loaded cells. Scale bars = 20 μm. Statistical analysis was performed using a Mann-Whitney non-parametric test (*P < 0.05, ^n.s^P = not significant). Summary statistics for boxplots: centre = median; bounds of box = IQR 25th and 75th percentile; whiskers = minimum and maximum within 1.5 IQR.

Notably, expression of MT-ND1 and MT-ATP6, genes (which encode for subunits of mitochondrial complexes I and V, respectively), was significantly lower following α-syn-treatment than in the control condition. Our live-cell tracking, therefore, aligns with previous work showing an impact on mitochondrial complexes I and V with the addition of aggregated α-syn.^34,35^ To understand if the change in mitochondrially encoded expression led to a functional impact on the mitochondria, a TMRM assay was performed on α-syn-treated and control cells (Fig. 5e). This showed a significant decrease in MMP in α-syn-treated cells compared to control cells.

These results suggest that the presence of aggregated α-syn decreases expression of mitochondrially encoded genes, such as MT-ND1 and MT-ATP6, in neurons, and this is concurrent with a decrease in MMP. However, we did not find a relationship between the proximity of aggregates to mitochondria and the effect on gene expression. Therefore, the effects of α-syn aggregates on mitochondrial gene expression may not be due to a direct interaction of the oligomers with the mitochondria themselves. Rather, the effect on mitochondria may be indirect, for example, as a downstream effect of α-syn toxicity in other regions of the cell.

## Discussion

In this work, using a precise and minimally invasive single-cell nanobiopsy technique, we extracted individual mitochondrion from live neurons with high spatial and temporal resolution. We demonstrate that single mitochondria can be extracted from neurons using the nanotweezer without affecting neuronal viability. The maintenance of MMP following mitochondrial extraction highlights the potential of the technique for mitochondrial transplantation, as has been achieved with other single-cell sampling approaches.^18,19^

Our development of methods for single-mitochondrial analysis demonstrates that information on mtDNA copy number, mtDNA mutations, and gene expression can be obtained from a single mitochondrion. We show that it is possible to analyse the RNA content of a single mitochondrion, opening opportunities for highly localised mitochondrial gene expression tracking and linking single organelle molecular changes with cellular function.

By dynamically measuring mitochondrial gene expression in the same cells over time, we tracked neuronal responses to the presence of α-syn aggregates in single cells. We revealed that these aggregates can lead to a downregulation of genes encoding subunits of mitochondrial complexes I and V. Given the dynamic nature of gene expression within cells, our approach enables the tracking of mitochondrial gene expression of a cell from its ground state to its treated state, unmasking the inherent heterogeneity and variability of cells and providing a more accurate depiction of single-cell responses. Interestingly, we find no correlation between the downregulation of MT-ND1 and MT-ATP6 at the single-cell level and the proximity of α-syn aggregates to individual mitochondria. This suggests that mitochondrial dysfunction may result from interactions with α-syn species that precede aggregate formation.

Overall, we demonstrate a powerful tool for extracting and analysing individual mitochondria in living neurons. A significant advantage of the method outlined here is the exceptional precision of the nanotweezer for selecting a mitochondrion of interest within a specific region of a cell, combined with minimal impact on the cell, enabling live single-cell measurements.

Current limitations of the technique include the number of mitochondrially encoded genes that can be analysed from a mitochondrion and the analysis of low-abundance transcripts. Improved preservation of nucleic acid content within each mitochondrion, combined with the ongoing development of highly sensitive sequencing methods, will further increase the power of the technique. We anticipate that the nanotweezer will become a valuable technique for studying the mitochondrial landscape of living cells.

## Methods

### Primary neuron culture

Primary hippocampal and cortical neuronal cultures were isolated from E18 Sprague Dawley rats. Cells were plated at a density of 25,000 cells/cm^2^ on 35 mm glass-bottom dishes (Greiner Bio-One) or gridded 35 mm glass-bottom dishes (Ibidi). Dishes were pre-coated with 0.25 mg/mL poly-L-lysine hydrobromide (Sigma-Aldrich) overnight. Cells were attached overnight in Minimum Essential Medium (Gibco) supplemented with 10% horse serum (Gibco), 1 mM pyruvic acid (Sigma-Aldrich), and 0.6% glucose solution (Sigma-Aldrich). Cells were then cultured in Neurobasal (Gibco) supplemented with 2% B27 (Gibco), 1% Glutamax (Gibco), 0.6% glucose (Sigma-Aldrich), and 1% Penicillin-Streptomycin (Gibco) at 37 °C with 5% CO2. Half media changes were performed every 3-4 days. Cells were transfected by lipofection 2 days before experiments using Lipofectamine 2000 (Life Technologies) and the fluorescent mitochondrial reporter DsRed2-Mito-7. For TMRM analysis, cells were incubated with 20 nM TMRM (Life Technologies) in cell maintenance media for 30 min at 37 °C and then washed in PBS and replaced with fresh media. DsRed2-Mito-7 was a gift from Michael Davidson (Addgene plasmid # 55838 ; http://n2t.net/addgene:55838 ; RRID:Addgene_55838).

### Immunocytochemistry

Immunocytochemistry was used to determine the spatial proximity of aggregates with mitochondria. Cells were then permeabilised with 0.1% Triton-X-100 (Thermo Scientific) in PBS for 15 min. Cells were washed in PBS and then blocked for 1 h at room temperature in 5% Bovine Serum Albumin (BSA) (Thermo Scientific) with 5% goat serum (Thermo Scientific) in PBS. anti-MAP2 (118 004, Synaptic Systems), anti-Tomm20 (ab56783, Abcam), and anti-alpha-synuclein aggregate (MJFR-14-6-4-2, Abcam) primary antibodies were diluted to 1:500, 1:100, and 1:5000, respectively, in blocking buffer and added to the samples overnight at 4 °C. The next day, samples were washed in PBS, and Alexa Fluor™ 647 goat anti-guinea pig (#A-21450, Invitrogen), Alexa Fluor 488 goat anti-mouse (#A-11001, Invitrogen) and Alexa Fluor 568 goat anti-rabbit (#A-11011, Invitrogen) secondary antibodies, each diluted 1:1000 in blocking buffer, were added to the samples for 2-3 h. Cells were washed in PBS and then mounted using ProLong™ Gold Antifade Mountant with DAPI (Invitrogen). Imaging was performed on a Zeiss Axio Observer widefield microscope.

### α-synuclein preparation and characterisation

N-acetylated α-synuclein monomers were expressed using recombinant expression in Escherichia coli. The concentration was estimated from the absorbance at 275 nm by using a molar extinction coefficient of 5600 M^-1^ꞏcm^-1^. The aggregation conditions were based on an established protocol.^34,40,41^ Briefly, 800 μl of α-syn monomers (70 μM, 25 mM Tris, 100 mM NaCl, pH 7.4) were ultracentrifuged for 1 h at 90,000 rpm and 4 °C to remove insoluble aggregates. The top 600 μl of the supernatant was collected to collect the monomers, while the remaining volume was discarded. 400 μl of this monomer sample was mixed with 0.01% NaN_3_ to prevent bacterial growth during aggregation. The mixture was then aggregated by incubating at a temperature of 37 °C with shaking at a constant speed of 200 rpm for 24 h.

For the production of fluorescently labelled α-synuclein aggregates, a fluorescently labelled aptamer specific for α-synuclein was used (5’-GGTGCGGGACTAGTGGGTGTGTTTTTT-ATTO647-3’, Integrated DNA Technologies). Aggregated α-synuclein was incubated with the aptamer for 1 h. The mixture was then washed with PBS and filtered 5 times using Amicon Ultra 100 kDa centrifugal filters to remove unbound aptamer and monomer.

α-Synuclein samples were characterised by AFM. α-Synuclein samples were placed on mica sheets and dried with a nitrogen gun. AFM imaging was performed using an Asylum MFP-3D microscope in tapping mode. Nanosensors PPP-FMR tips (res ∼75 kHz, nominal tip radius: 7 nm, nominal spring constant: 2 N/m) were used and tuned to a target tapping amplitude of 1 V. Scans were sized 90 μm^2^, 256-512 points per line, with a scan rate of 0.5 Hz.

### Nanobiopsy

Nanotweezers were fabricated, and single mitochondrial nanobiopsies were performed using the nanotweezer, as reported previously.^25^ Briefly, mtdsRed-transfected neurons were mounted on an inverted optical microscope (IX71 or IX83, Olympus). Copper wires were inserted into each chamber of the nanotweezer, and the opposite ends were connected to a function generator (TG2000, TTi). The nanotweezer was mounted above the cells and manipulated using a micromanipulator. (PatchStar, Scientifica). Guided by bright-field illumination and fluorescence microscopy, the nanotweezer was manually inserted into the neurite of a neuron adjacent to a labelled mitochondrion. An AC voltage (1 MHz, 15 V peak-to-peak) was applied using the function generator to trap a single mitochondrion dielectrophoretically. The nanotweezer was then retracted from the neuron whilst maintaining the DEP force, and the trapped mitochondrion was transferred to a PCR tube by pressing the tip into the tube containing 4 μl lysis buffer (100 mM Tris-HCl [pH 8], 150 mM NaCl, 20 mM EDTA, 0.2% SDS) in nuclease-free water. For mtRNA analysis, the lysis buffer also contained 1 unit RNAase inhibitor (#N8080119, Applied Biosystems).

### Single mitochondrion lysis and purification

1 μl 0.05 mg Proteinase K was added to the lysis buffer containing the isolated mitochondrion, and the sample was lysed at 55 °C for 30 min. For mtDNA analysis, single mitochondrion lysates were purified using AMPure XP Bead-based reagent (Beckman Coulter) with a modified protocol. The reagent was vortexed before use to ensure homogeneous bead distribution. 7 μl beads were added to 5 μl mitochondrial lysate (1.4x) in a 0.5 mL tube, mixed by gentle pipetting 10 times and then incubated for 5 min at room temperature to allow for the DNA to bind to the beads. The tube was placed on a magnetic rack (#S1509S, NEB) for at least 2 minutes to allow the beads to separate from contaminants and lysis buffer components. The supernatant was removed, leaving approximately 2 μl in the tube. Whilst still bound to the magnet, the beads were washed twice with 50 μl fresh 70% ethanol in nuclease-free water. Full removal of the supernatant was performed at each wash step to further remove contaminants and lysis buffer components. Immediately after the last wash step, to minimise bead drying, 5-10 μl nuclease-free water was added to the beads and mixed thoroughly, away from the magnetic rack. Beads were incubated for 2 min to elute DNA. The tube was then placed back on the magnetic rack for 2 min to ensure the solution was fully transparent, and the entire eluate was transferred to a PCR tube.

For mtRNA analysis, two types of surface-functionalised magnetic beads were generated to allow for specific binding to rat mtRNAs. DynabeadsTM MyOneTM Carboxylic Acid (#65011, Invitrogen) 1 μm magnetic beads were functionalised with oligonucleotides complementary to mtRNAs via a carbodiimide coupling reaction following the manufacturer’s protocol. An oligonucleotide comprising a repeated sequence of 25 thymine nucleotides for binding to polyadenylated RNAs (5’-H_2_N-C_6_-PO_4_^—^ oligo(T)_25_-3’, Integrated DNA Technologies), and a second oligonucleotide comprising a sequence complementary to 12S rRNA (5’-H2N-C6-PO_4_-oligo(TAATAAGGTTCGTGTGAAAGGTCAT)-3’, Integrated DNA Technologies) were incorporated into the reaction.

### qPCR primers and probes

All primers and probes used for qPCR are listed in Table S1. Primers and probes were designed using Primer-BLAST (https://www.ncbi.nlm.nih.gov/tools/primer-blast/), PrimerQuest (https://eu.idtdna.com/PrimerQuest/) or ordered commercially (Integrated DNA Technologies). For multiplexed qPCR, primer/probe compatibility was validated using the OligoAnalyzer™ Tool (https://eu.idtdna.com/pages/tools/oligoanalyzer/) and the Eurofins Genomics Oligo Analysis Tool (https://eurofinsgenomics.eu/en/ecom/tools/oligo-analysis/).

### qPCR

For mtRNA analysis, purified mtRNA samples were converted to cDNA using the iScript cDNA Synthesis Kit (Bio-Rad) following the manufacturer’s instructions. qPCR reactions were performed using a StepOnePlus Real-Time 96-well PCR system (Applied Biosystems) in MicroAmp™ optical 0.1 mL 96-well plates (Applied Biosystems) or using a QuantStudio 6 Flex system in MicroAmp™ optical 384-well plates (Applied Biosystems). For initial validation experiments (Fig. 2), singleplex reactions were performed on cDNA samples using SsoAdvanced Universal SYBR Green Supermix (Bio-Rad) over 40 cycles, with thermocycling and reaction conditions set according to the manufacturer’s protocol. A melt curve stage was run after the amplification protocol by increasing the temperature at a rate of 0.5 °C/s from 60 °C to 90 °C. All other experiments were performed as duplex reactions using PrimeTime™ Gene Expression Master Mix (Integrated DNA Technologies). A 2:1 primer:probe ratio (500 nM:250 nM) was incorporated with thermocycling over 40-45 cycles according to the manufacturer’s protocol. Copy numbers for all targets were determined by absolute quantification using a standard curve of known copy numbers. Standard curves were run on the same plate where possible. Otherwise, the most recent standard curve was used, and the threshold for sample target amplification was set to the standard curve’s threshold.

### Sequencing

A 558 bp region of the MT-RNR2 gene of mtDNA was amplified using high-fidelity PCR (Q5 High-Fidelity 2x Master Mix, New England Biolabs) according to the manufacturer’s protocol. The PCR product was purified using the GeneJET Gel Extraction and DNA Cleanup Micro Kit (Thermo Scientific), and the concentration was determined using a Qubit Fluorometer (Invitrogen). 150 fmol of DNA was sequenced using the MinION device (Oxford Nanopore Technologies) with the MinION Flow Cell (R10.4.1) and the Ligation Sequencing Kit V14 (SQK-LSK114) according to the manufacturer’s protocol. The MinKNOW software (Oxford Nanopore Technologies) was used to collect the raw data and convert it into basecalled reads. Post-basecalling analysis was performed using the EPI2ME platform (Metrichor Ltd.) for sequence alignment.

### Analysis

Images were analysed using Fiji. Sequencing analysis was performed using Integrated Genomics Viewer (IGV) and samtools.^42,43^ Data processing was performed using MATLAB 2020 or Origin 2020. Normality and statistical tests were performed using GraphPad Prism 8. The Shapiro-Wilk test was used to test for normality before all statistical tests. A non-parametric test was used if the data failed the normality test. p < 0.05 was considered statistically significant.

## Authors and affiliations

Annie Sahota − Department of Chemistry, Imperial College London, Molecular Science Research Hub, London W12 0BZ, United Kingdom; orcid.org/0000-0003-2923-1550

Binoy Paulose Nadappuram − Department of Chemistry, Imperial College London, Molecular Science Research Hub, London W12 0BZ, United Kingdom; Department of Pure and Applied Chemistry, University of Strathclyde, Glasgow G1 1BX, United Kingdom; orcid.org/0000-0002-1386-8357

Siân C. Allerton − Department of Chemistry, Imperial College London, Molecular Science Research Hub, London W12 0BZ, United Kingdom.

Flavie Lesept − Department of Neuroscience, Physiology and Pharmacology, University College London, London WC1E 6BT, United Kingdom.

Jack H. Howden − Department of Neuroscience, Physiology and Pharmacology, University College London, London WC1E 6BT, United Kingdom.

Suzanne Claxton − Kinases and Brain Development Lab, The Francis Crick Institute, London NW1 1AT, United Kingdom.

Yaxian Liu − Department of Chemistry, Imperial College London, Molecular Science Research Hub, London W12 0BZ, United Kingdom.

Francesco A. Aprile − Department of Chemistry, Imperial College London, Molecular Science Research Hub, London W12 0BZ, United Kingdom; orcid.org/0000-0002-5040-4420

Josef T. Kittler − Department of Neuroscience, Physiology and Pharmacology, University College London, London WC1E 6BT, United Kingdom.

## Contributions

A.S., B.P.N., A.P.I., J.B.E., and M.J.D. conceived and designed experiments. A.S. performed the experiments. S.C.A. and Y.L. contributed to the optimisation of the α-synuclein experiments. A.S. wrote the manuscript. F.L., J.H.H., and S.C. isolated the primary neurons. F.A.A. contributed to the α-synuclein samples. J.T.K. contributed to the cell samples and experiments. A.P.I., J.B.E., and M.J.D. were responsible for the conception and supervision of the project. All authors have approved the final version of the manuscript.

## Acknowledgments

We thank the Facility for Imaging by Light Microscopy and the Electron Microscopy Facility at Imperial College London. We thank Dr Victoria Bemmer at the microscopy facility at Imperial College London for the AFM images. We thank the Biological Research Facility at the Francis Crick Institute for animal support and Dr Sila Ultanir for providing the primary neuron cultures. We also thank Dr Xiangyu Teng and Dr Liming Ying for providing initial α-synuclein samples and Dr Caroline Koch for guidance on the MinION sequencing.

A.S. and S.C.A. were supported by a scholarship from the EPSRC CDT in Chemical Biology (EP/S023518/1). A.S. also acknowledges support from UKRI/Wellcome grant EP/T022000/1−PoLNET3. A.P.I. and J.B.E. acknowledge support from EPSRC grant EP/V049070/1. F.A.A. acknowledges support from UKRI (Future Leaders Fellowship MR/S033947/1 and MR/Y003616/1). This project has also received funding from the European Research Council (ERC) under the European Union’s Horizon 2020 research and innovation program (grant agreement nos. 724300 and 875525). J.T.K. is supported by Wellcome Trust Senior Investigator (22519/Z/21/Z) and Collaborator Awards (223202/Z/21/Z). M.J.D. was supported by the Francis Crick Institute, which receives its core funding from Cancer Research U.K. (CC2206), the U.K. Medical Research Council (CC2206), and the Wellcome Trust (CC2206).

