## Supporting Information for "Tracking gene expression of single mitochondria in live neurons using nanotweezers"

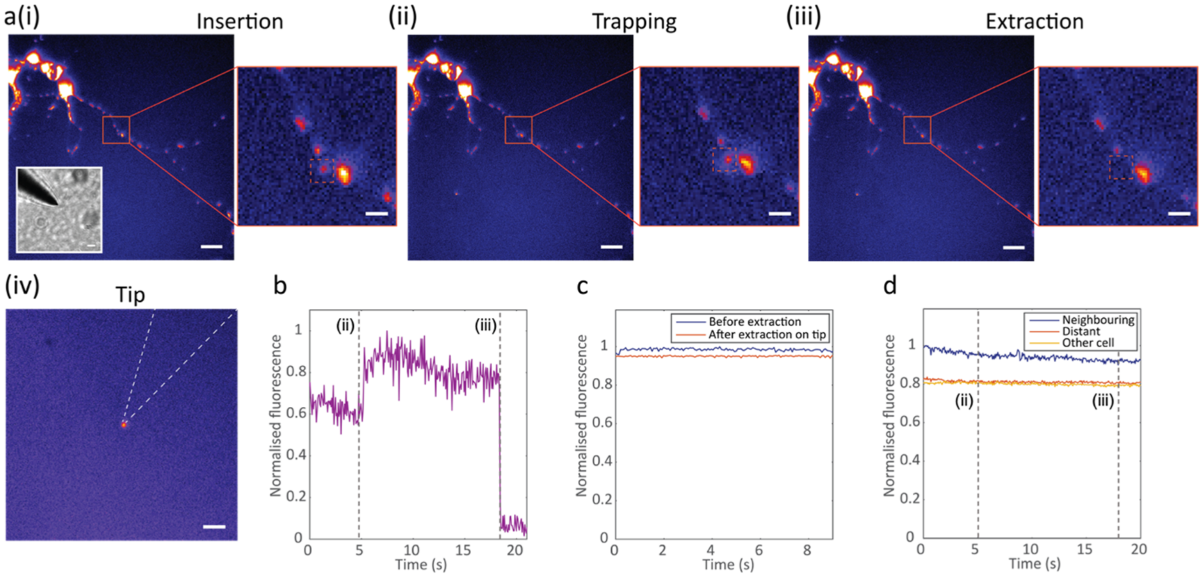


**Fig. S1| Mitochondrial viability during single mitochondrion isolation.** a) Mitochondria were labelled with TMRM to monitor active mitochondrial membrane potentials. The nanotweezer was inserted into a neuron next to a labelled mitochondrion (i), DEP was applied to trap the mitochondrion, showing no loss or change in mitochondrial membrane potential (ii), the nanotweezer was removed from the cell to extract the mitochondrion (iii) and the tip of the nanotweezer was visualised outside the cell to confirm extraction of a viable mitochondrion (iv). Scale bars = 20 μm, zoomed scale bars = 10 μm, tip (iv) scale bar = 10 μm. b) Fluorescence-time trace of the nanotweezer tip during the trapping procedure, showing the time point where DEP was applied for mitochondrion trapping at the tip (ii) and when the nanotweezer was removed from the cell, leading to a loss of fluorescence from removal of the mitochondrion (iii). c) Fluorescence of the targeted mitochondrion before (blue) and after (orange) nanotweezer extraction, showing no loss of mitochondrion membrane potential. Fluorescence was normalised to the largest measured value. d) Fluorescence of non-targeted neighbouring mitochondrion (blue), distant mitochondrion from the same cell (orange) and mitochondrion from a different cell (yellow) during the extraction procedure, with the timepoints of DEP application (ii) and extraction of the targeted mitochondrion (iii) shown by the dashed lines. No significant change in fluorescence and membrane potential of other mitochondria was observed during the isolation procedure. Fluorescence was normalised against the largest fluorescence value measured across mitochondria.

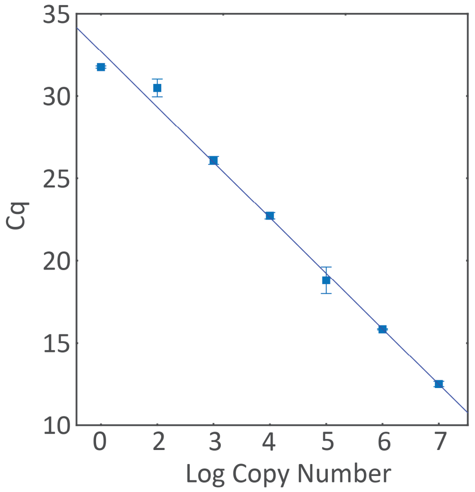


**Fig. S2| Standard curve for absolute quantification of mtDNA copy number (MT-CO1).**

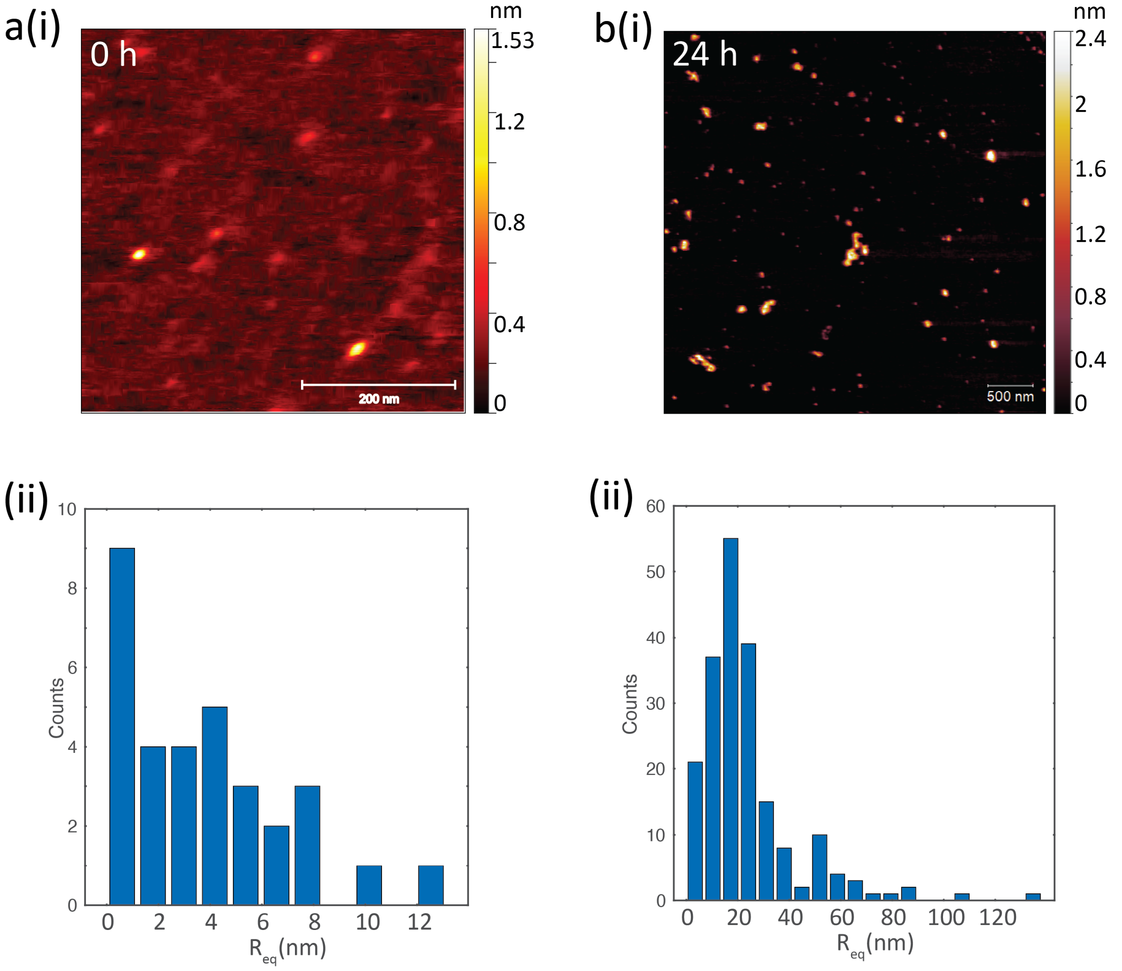


**Fig. S3| Characterisation of α-synuclein aggregation.** a) AFM images (i) and histograms of the equivalent disc radius ($R_{eq}$) (ii) of α-synuclein monomers before aggregation. b) AFM images (i) and histograms of the equivalent disc radius ($R_{eq}$) (ii) of α-synuclein following aggregation for 24 h.

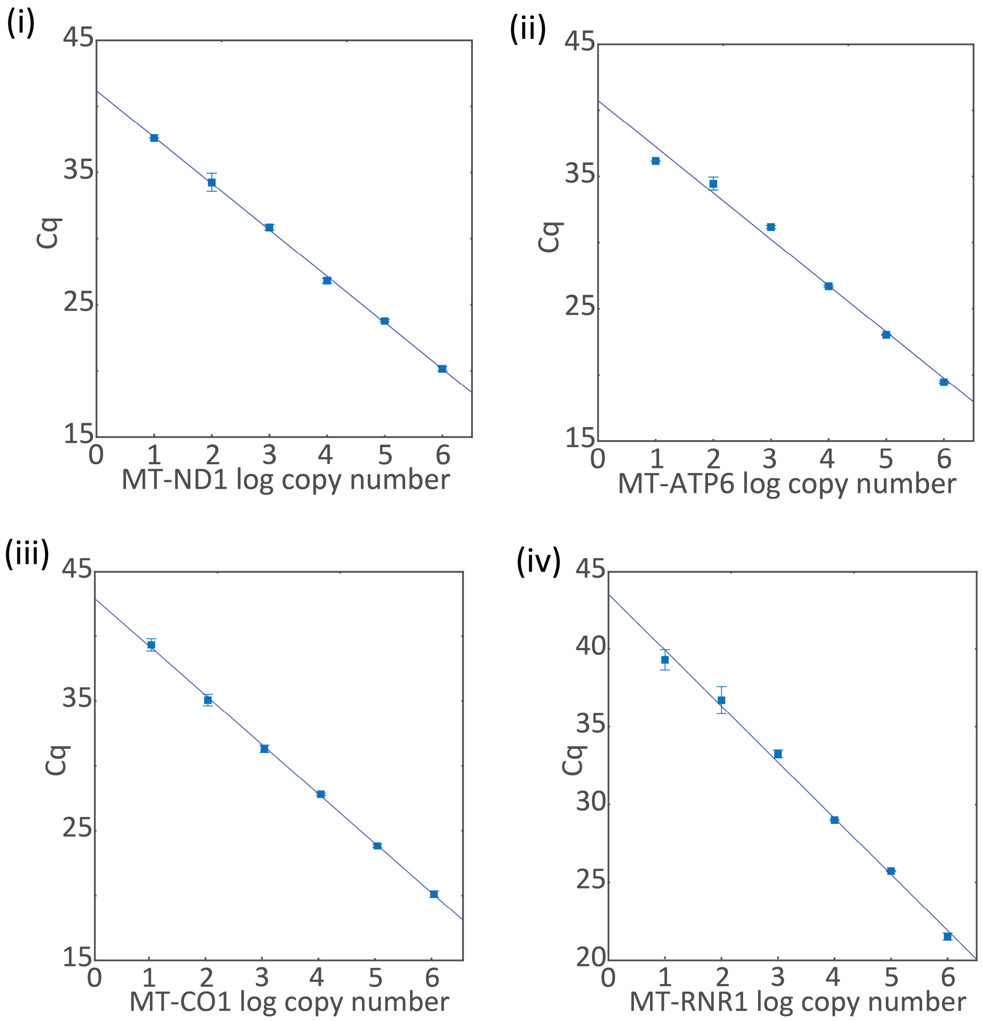


**Fig. S4| Standard curves for absolute quantification of single mitochondrion gene expression.** Representative qPCR standard curves for MT-ND1 (i), MT-ATP6 (ii), MT-CO1 (iii) and MT-RNR1 (iv).

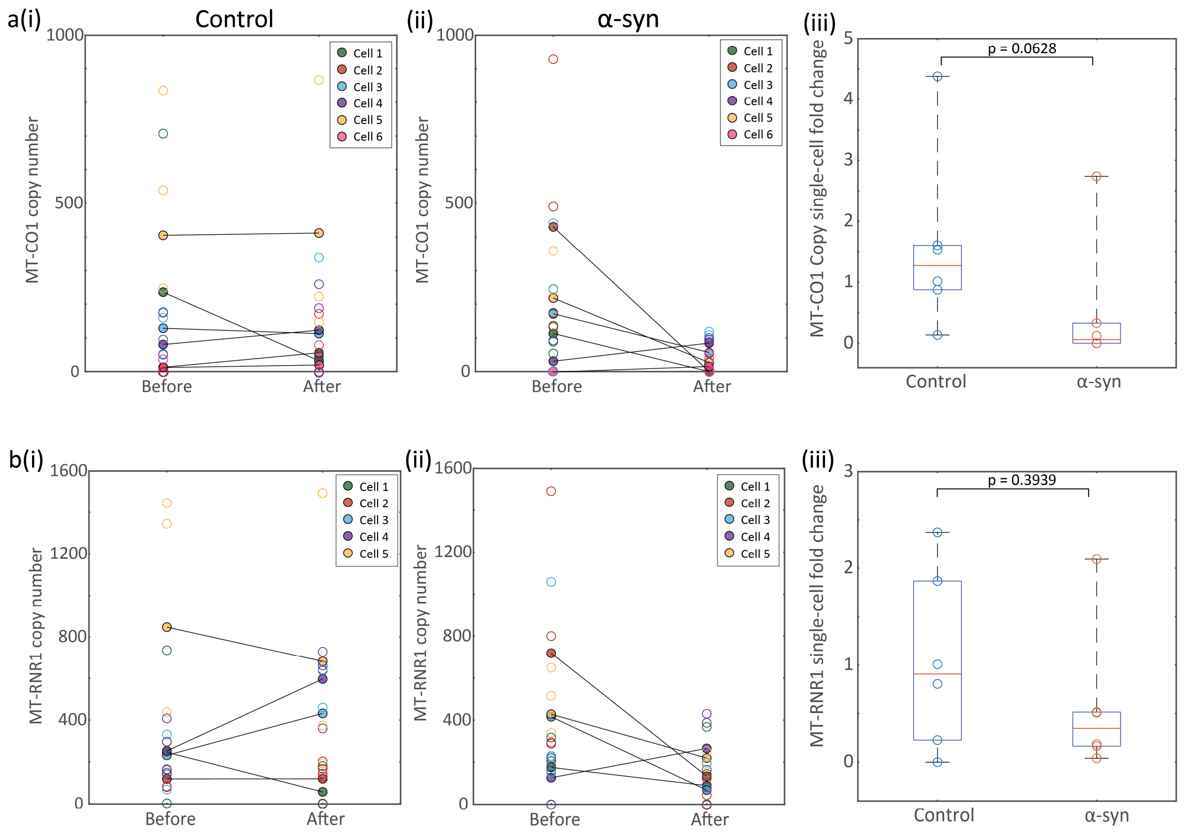


**Fig. S5|** ***Single-cell tracking of mitochondrially-encoded gene expression with application of aggregated α-synuclein.*** Expression of CO1 tracked in control cells (i), expression of MT-CO1 tracked in α-synuclein-treated neurons (ii), and (iii) the fold change of MT-CO1 measured in the same single cells in control and α-synuclein-treated neurons (n=6 cells). d) Expression of MT-RNR1 tracked in control cells (i), expression of MT-RNR1 tracked in α-synuclein-treated neurons (ii), and (iii) the fold change of MT-RNR1 measured in the same single cells in control and α-synuclein-treated neurons (n=6 cells). Statistical analysis was performed using a Mann-Whitney non-parametric test. Each filled circle in the tracked plots represents the mean copy number of each cell; the unfilled circles represent the individual copy numbers from each mitochondrion, and the black lines link the mean copy numbers in the same cell before and after treatment. Each unfilled circle in the fold change plots represent the fold change between the mean mitochondrial copy numbers in the same cell before and after treatment.

**Table S1| List of primer and probe sequences used in qPCR.**

| Target | Forward primer | Reverse primer | Probe |
|---|---|---|---|
| MT-CO1 (dye-based) | ATCAAATGATCCCCCGCCAT | GTACGATCCCTGTTAGGCCC | N/A |
| MT-CO1 | ATCAAATGATCCCCCGCCAT | GTACGATCCCTGTTAGGCCC | TGAGCCTTAGGGTTTATCTTCTTATTCACAGT |
| MT-RNR1 | TACCGCCATCTTCAGCAAAC | TTCCGCTTCATTGGCTACAC | AGGCACTAAAGTAAGCACAAGAACAAACA |
| MT-ND1 | TCTTACGAAGTCACAATAGCCA | AGTCAGATATGTTCTTGTGTAGTGA | CCGTCCTCCTAATAAGCGGCTCCTT |
| MT-ATP6 | ATGCATCTAATCGGAGGAGCTA | TTAAGGCTACGGCAAATTCAAG | AGACATCAGCCCACCAACCGCTACA |

All primers/probes were used in probe-based qPCR unless stated as dye-based.
